# Bringing Attention to Education: Revisiting the Neural Mechanisms of Selective Attention in Naturalistic Learning

**DOI:** 10.64898/2026.08.26.747092

**Authors:** Adi Korisky, Radhika S Gosavi, Shelly Whittet, Elizabeth Y Toomarian, Vani Dewan, Blair Kaneshiro, Bruce D McCandliss

**Affiliations:** Graduate School of Education, Stanford University, Stanford, CA, USA; Synapse School, Menlo Park, CA, USA

**Author notes:** Correspondence: Adi Korisky.

**Keywords:** Auditory Attention, Speech processing, EEG, Neuroeducation, Research-practice partnerships

## Abstract

In the context of education, attention can be considered the gateway for learning. Cortical attention mechanisms may help explain, in a precise and mechanistic manner, crucial educational modulators that are essential to effective learning and relevant to constructs that are essential to developing learners. Bringing cortical attention mechanisms into dialogue with educational practice, however, raises a fundamental challenge: which of the multitude of proposed cortical mechanisms that have been operationalized within the context of the lab are most relevant to classroom learning? Here we seek to address this challenge via attentional manipulation of EEG signals produced by middle school students as they experience an authentic novel instructional session within their school. First, we work with practitioners to co-construct naturalistic auditory and visual educational streams of information, and then manipulate student audio-visual learning priorities to examine the effects on the temporal dynamics of EEG signals linked to their teacher’s voice. We conduct temporal response function analyses across whole-topography components to help differentiate the relevance of early (e.g. early acting sensory modulation) versus late (biased competition or conflict resolution) attention mechanisms. Significant attentional modulation of speech processing emerged at only 170 ms, lending support to late but not early selection mechanisms. Moreover, individual differences in such cortical attention effects proved to be relevant to learning abilities and to teacher impressions of how well each student was able to engage during real classes. These approaches to naturalistic EEG within schools may guide selection of candidate mechanisms of attention within future research that seeks to leverage mechanistic insights from cognitive neuroscience to improve educational practice.

## 1. Introduction

Throughout early development, children learn by listening to more knowledgeable others. In early childhood, learning occurs in one-on-one interactions in which attention is supported by immediate feedback and adaptive communication, allowing the learner to develop (Vygotsky, 2011). Formal schooling, however, places fundamentally different attentional demands on learners. A single teacher must communicate with dozens of students simultaneously, requiring each student to independently regulate their attention, identify relevant information within ongoing instruction, and resist competing internal and external distractions. Studies have shown that the ability to navigate these attentional demands in the classroom is critical for successful learning and broader life outcomes, with difficulties in attentional regulation associated with lower academic achievement, reduced academic motivation, and an increased risk of school failure (Eccles, 2004; Polderman et al., 2010; Treviño et al., 2021). Yet, despite the central role of attention in educational success, the neural mechanisms that enable students to successfully regulate their attention during naturalistic learning remain poorly understood.

Growing efforts to study attention in ecologically valid contexts have sought to better capture the attentional demands of everyday life. Initially, these efforts focused on the nature of the presented stimuli. One line of research has focused on increasing the ecological validity of the target auditory stream, moving beyond isolated, artificial stimuli presented in repeated-trial designs toward continuous speech and examining how attention shapes its neural processing in the presence of competing auditory streams (Ding & Simon, 2012, 2014; Horton et al., 2013; Kaufman & Zion Golumbic, 2023; Kerlin et al., 2010; Li et al., 2022; Mesgarani & Chang, 2012; O’Sullivan et al., 2015; Zion Golumbic et al., 2013). This approach not only increases the ecological validity of experimental findings but also enables a better exploration of the temporal dynamics of attention in complex listening scenarios. Across these studies, a relatively consistent finding is that attentional modulation is most reliably observed after approximately 100 ms and centered on the P1–N1–P2 complex. However, the specific component within this complex at which attentional modulation emerges, as well as the precise latency and magnitude of these effects, varies considerably across the literature (Fritz et al., 2007).

A second wave in the movement toward studying attention under more naturalistic conditions extended beyond the auditory modality to investigate attentional deployment in multimodal environments that better reflect the complexity of everyday life. Motivated by the observation that speech in real-world settings is rarely encountered in isolation but is typically accompanied by competing visual information, researchers adapted classic audio–visual selective attention paradigms (Gomez-Ramirez et al., 2011; Korisky et al., 2026; Lakatos et al., 2008; Schroeder & Lakatos, 2009) to examine the neural mechanisms underlying shifts of attention toward or away from continuous natural speech while participants simultaneously processed competing visual stimuli. These cross-modal studies highlight the role of cognitive effort in shaping auditory processing, showing that attentional modulation emerges selectively within the P2 window, but only when speech comprehension or the processing context places high demands on the listener (Kong et al., 2014; Vanthornhout et al., 2019; Xie et al., 2023). Whereas auditory-only paradigms place attention in direct competition within a single sensory domain, cross-modal paradigms reduce this competition, showing that under low cognitive demands, participants continue to track the speech envelope even while attending to another sensory stream.

The paradigms above each capture at least one potential aspect of attention, each with its own cortical time dynamics as they modulate continuous speech. This raises a vexing but critical question in the progression toward studying attention within naturalistic learning environments: which neural timescales and modulatory patterns are ultimately most relevant when speech perception is embedded within a complex, multi-modal learning environment?

The context of education represents a critical arena for validating these laboratory-derived attention effects and adjudicating which are most relevant to supporting learning. During classroom learning, students must track instructional speech not merely to decode acoustic features and process linguistic intelligibility, but to actively drive high-level cognitive operations, such as constructing mental schemas, organizing novel information, and integrating incoming content with prior knowledge (Mayer, 2014, 2024). These high-level processes must execute continuously within a multisensory environment where auditory and visual inputs dynamically converge or compete for finite attentional resources (for example, when a student listens to their teacher while simultaneously extracting meaning from complex diagrams on a worksheet or text on a presentation slide). Given this complexity, it remains unclear whether attentional mechanisms characterized in simplified laboratory paradigms adequately capture the processes that support learning in authentic classrooms. Consistent with this concern, a growing body of work has shown that traditional laboratory measures of attention often fail to predict real-world educational performance (Faraone et al., 2021; Parsons et al., 2017; Slattery et al., 2022), underscoring the need for better translational approaches moving forward (Janssen et al., 2021; van Atteveldt et al., 2020).

In the present study, we extend the progression toward ecologically valid attention research by investigating how discrete temporal stages of neural speech tracking, early sensory versus late top-down processes, adapt to the cross-modal cognitive demands of classroom learning in adolescents. We adopt an individual-level perspective by examining whether students’ ability to flexibly modulate auditory attention in response to educational demands predicts speech processing, learning outcomes, and academic performance. To simulate learning, we co-developed a naturalistic audiovisual selective attention paradigm with an experienced classroom teacher that captures key features of classroom learning. Using school-based electroencephalography (EEG), fifth- and sixth-grade students engaged in curriculum-aligned lessons while shifting their attention between their teacher’s spoken instruction and a concurrent visual learning task. A linear temporal response function (TRF) approach was applied to quantify auditory envelope tracking in the presence of visual distraction, incorporating complementary encoding (forward) and decoding (backward) models. We hypothesized that the cognitive demands inherent to learning would elicit late-stage attentional modulation (∼200 ms), even in the absence of competing auditory demands, while leaving early sensory encoding (∼100 ms) relatively unaffected. Importantly, we tested whether these neural mechanisms translate beyond the experimental task, predicting individual differences in both learning outcomes and real-world classroom attention.

## 2. Methods

### 2.1 Participants

Forty-seven 5^th^-6^th^ grade students participated in the study. Each participant completed a single EEG session in a school-based EEG lab (Toomarian et al., 2024). Participants were recruited from the school, and sessions were conducted during the school day. From this cohort, two datasets were excluded due to technical issues during the sessions, and two additional datasets were excluded because participants did not complete the full session. In additional four datasets were excluded due to excessive motor artifacts, affecting more than 60% of the recording. Thus, the presented analysis shows data from 39 participants in total (age range: 10.2–12.3 years, M = 11.31, SD 0.60; 21 girls). All students who provided verbal assent and whose parents provided written informed consent were allowed to participate in the study, regardless of psychological or medical diagnoses. Demographic data and information on participants’ diagnoses, medical history, and current prescribed stimulant medication were collected from parents and are reported in Table S1. The study was approved by the Stanford University Institutional Review Board, and each participant received a small token gift as compensation for participation.

### 2.2 Co-Creation Process with the Partner Teacher

To develop an ecologically valid paradigm that captures attentional processes during real-world classroom learning, the experimental task was created through an iterative four-phase co-creation process between the research team and an experienced classroom teacher (Figure 1). The partner teacher had over a decade of teaching experience and worked directly with the students who participated in the study, providing extensive knowledge of their curriculum, learning needs, and classroom dynamics.

**Figure 1:**
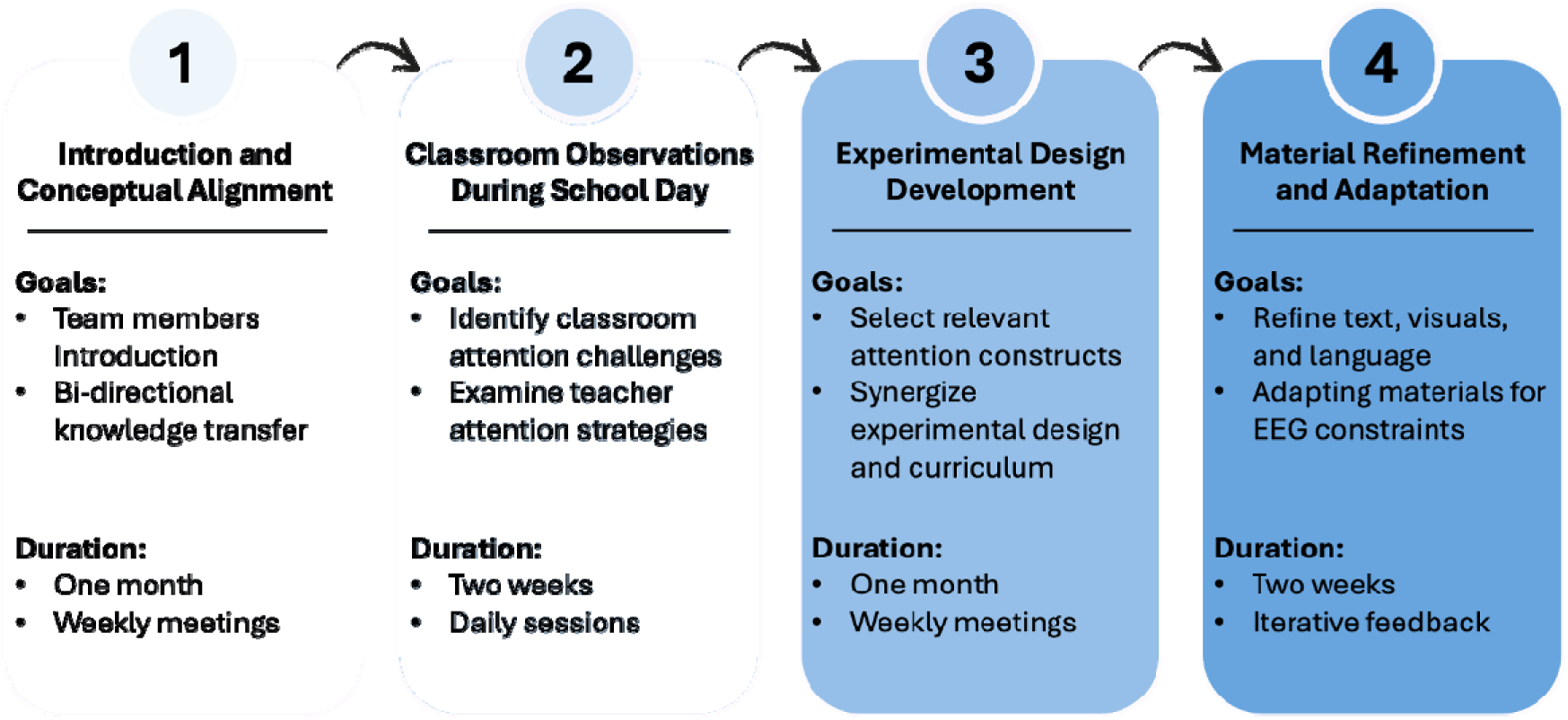
Co-creation process underlying the development of the experimental paradigm. The experimental paradigm was developed through a four-phase research–practice partnership between the research team and an experienced classroom teacher over approximately three months. The process included (1) conceptual alignment of neuroscientific and educational perspectives on attention, (2) classroom observations to identify authentic attentional demands and instructional strategies, (3) collaborative development of an EEG-compatible paradigm using curriculum-relevant materials while preserving key features of classroom learning, and (4) iterative refinement of the task, stimuli, and procedures to ensure pedagogical authenticity, age appropriateness, and experimental feasibility.

Based on previous models of research-practice partnerships (Gosavi et al., 2026; Korisky et al., 2024; Toomarian et al., 2024; Wang et al., 2025), the co-creation began with discussions to establish a shared understanding of attention from neuroscience and educational perspectives, followed by classroom observations to identify attentional demands and instructional strategies in authentic learning environments. These insights informed the development of an EEG-compatible paradigm using educationally meaningful content while preserving key features of classroom learning. The co-design process spanned approximately six months, with meetings between the research team and the teacher occurring, on average, once a week and increasing in frequency as the study launch approached.

### 2.3 Task Description

Our co-creation process yielded an audio–visual selective attention task comprising two conditions: Active Listening and Passive Hearing (Figure 2). During each experimental block, auditory and visual stimuli were always presented simultaneously, and participants were instructed to selectively attend to one stream while ignoring the other. To create an engaging and educationally meaningful context, the task was framed as a museum visit, in which students accompanied their teacher on guided museum tours and were asked to pay attention to their teacher’s voice (‘Active listening’) or to ignore it and perform a visual task (‘Passive hearing‘). The experimental context was aligned with the school’s annual curriculum theme, and all materials and procedures were iteratively refined with the teacher to ensure that the task was age-appropriate, pedagogically meaningful, and experimentally feasible. All photographers included in the experiment were selected because they were not part of the regular school curriculum.

**Figure 2:**
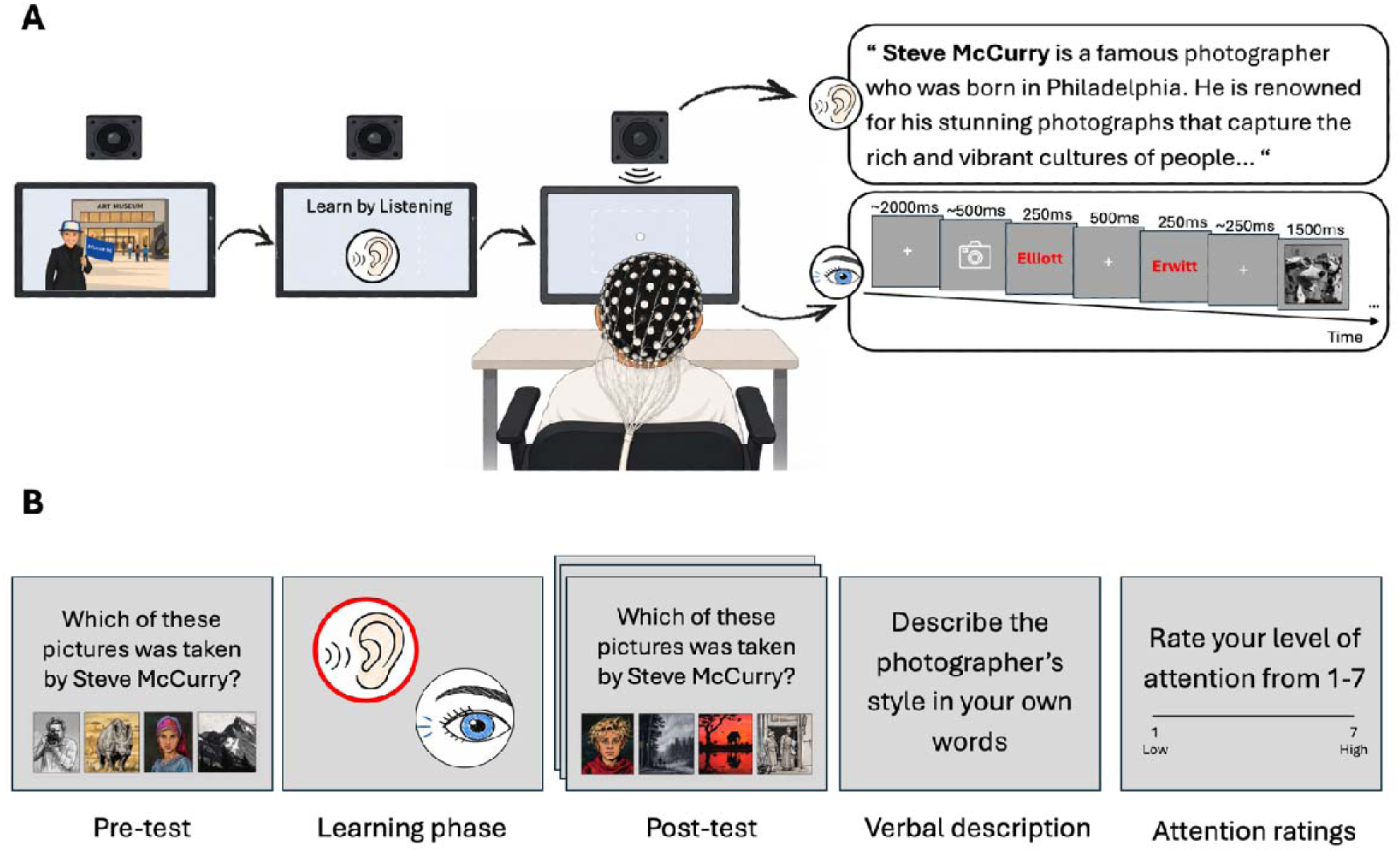
Experimental design and behavioral assessments. (A) Schematic illustration of the learning phase, which consists of an audio-visual selective attention task. Students listened to naturalistic educational content while EEG activity was recorded using a 128-channel EGI system. During each experimental block, students viewed a 2.5-minute-long movie containing simultaneously presented auditory and visual streams and were instructed either to listen actively to their teacher (“Learn by Listening”, Active listening condition) or to ignore the auditory stream and focus on the visual information (“Learn by Looking”, Passive hearing condition). Throughout the block, students were instructed to maintain visual fixation on the screen. The auditory stimulus was composed of continuous natural speech. Structure of the visual stimulus sequence. Each trial consisted of a fixation screen (∼2000 ms), a camera cue (∼500 ms), the photographer’s first name (250 ms), a fixation screen (500 ms), the photographer’s last name (250 ms), a second fixation screen (∼500 ms), and the corresponding image (1500 ms). (B) Schematic representation of one experimental block and associated behavioral measures. Across the experiment, participants completed six experimental blocks in total, consisting of three active-listening blocks (“Learn by Listening”) and three passive-listening blocks (“Learn by Looking”). Prior to each block, students completed a familiarity pre-test in which they identified photographers from image arrays. Following each block, students were asked to complete a post-test learning assessment composed of novel images (pictures matrix), provide a verbal description of the photographer’s style, and rate their attention and level of interest during the block.

*The auditory stimulus* simulated a short classroom lesson in which students learned about a photographer through teacher-led instruction. Each block consisted of approximately 2.2 minutes (M = 2.12, SD = 0.01) of natural speech recorded by the students’ teacher. During each block, the teacher described the life, artistic style, and photographic characteristics of one well-known photographer (e.g., Steve McCurry, Ansel Adams, and Dorothea Lange). All auditory stimuli were recorded by the teacher in their natural voice and speaking rhythm within the school-based laboratory using a professional microphone. The recordings were subsequently edited in Audacity (version 3.7; https://www.audacityteam.org/) to adjust sound intensity and onset of speech across recordings.

*The visual stimulus* imitated a visit to the museum and consisted of 32 image-name sequences, each combining a photographer’s name with one of their photographs, drawn from five different photographers (see Figure 2A for more details). From the 32 sequences, 12 corresponded to one target photographer and were highlighted in red to facilitate identification, whereas the remaining 20 image presentations consisted of distractor photographers (five repetitions per photographer). The photographer discussed in the simultaneous auditory stream never appeared in the concurrent visual stream, preventing overlap between the auditory and visual learning materials. Also, image presentations were pseudo-randomized such that target-photographer events did not appear consecutively more than twice in succession. In the current paper, we focus on the role of attention in shaping neural responses toward speech; thus, visual processing and related neural responses are beyond the scope of the current work.

Custom-made MATLAB scripts were used to generate the auditory and visual streams according to the criteria described above and combine them into unified MP4 files. All stimuli were presented in full-screen mode using the Psychtoolbox package in MATLAB (R2024b, version 24.2.0). To control for stimulus-specific effects, three such pre-recorded movies, including both auditory and visual stimuli combined, were presented twice to each participant. For each movie, participants completed the task once in the active listening condition and once in the passive hearing condition, resulting in a total of six experimental blocks. Although the same audio-visual material was repeated across conditions, block order and condition order were randomized across participants, with the constraint that the same movie or the same attention condition could not appear consecutively.

### 2.4 Experimental Procedure

In each session, participants were seated in a school-based EEG lab in a sound-attenuated room. Lighting was calibrated to be consistent across participants, and all participants were seated at a viewing distance of approximately 1 m from a computer screen. Auditory stimuli were delivered in a free-field manner through a loudspeaker positioned above the monitor to preserve a classroom-like listening environment. Participants were asked to maintain their gaze toward the screen throughout both attention conditions, and gaze direction was monitored by a member of the research team during the session. All participants were familiarized with the EEG net and informed about the goals of the study. Before beginning the experiment, students signed an assent form in addition to the written informed consent previously provided by their parents to confirm their willingness to participate in the session.

Participants were introduced to the task through the framing of a “museum visit,” in which their teacher guided them through a series of educational tours about famous photographers. Students were informed that their goal was to learn about the photography style of each photographer by listening (‘*Active listening’*) or by looking (‘*Passive hearing’*). In the *active-listening condition*, participants were instructed to focus on their teacher’s speech while maintaining visual fixation on the screen and ignoring the concurrent visual information. In the *passive-hearing condition*, participants were instructed to ignore the teacher’s ongoing speech and focus on the screen to learn from the visual cues about the photography style of a target photographer. In both conditions, participants did not perform any task during the learning phase, but were informed beforehand that their knowledge would be tested afterward.

A guided practice trial for each condition separately and the combined audio-visual block were conducted at the beginning of each session to ensure understanding of the task instructions. To assess learning outcomes, participants completed two assessments following each block: (1) a three-trial ‘*Pictures-matrix’* test, in which they identified novel photographs taken by the photographer introduced in the current block (‘*post-test*’, Figure 2B) and (2) a ‘*verbal description’* task, in which they were asked to describe the photographer’s visual style based on what they had learned during the experimental block. All test images were novel, had not been presented during the experimental blocks, and were not explicitly described in the auditory narratives. Thus, successful performance required participants to generalize learned stylistic features to new exemplars rather than rely on their memory solely. To rule out familiarity effects of the content that could impact learning, before each experimental block, participants completed a baseline familiarity assessment (‘*pre-test*’, Figure 2B) in which they reported whether they had previously heard of the target photographer and attempted to identify the photographer’s work from a four-alternative image matrix. At the end of each block, participants were asked to rate their level of attention (“*Please rate your level of attention during the last tour*”) and interest (“*Please rate how interesting this tour was*”) using a 7-point Likert scale (See Figure 2B for the full design).

The experiment included six blocks (i.e., ‘museum visits’), three for each condition. Short breaks were provided between blocks according to participants’ needs. Full sessions lasted approximately 55 minutes, including EEG setup, instructions, practice trials, experimental blocks, learning assessments, and between-block breaks.

### EEG Recording

Neural activity was recorded using a 128-sensor EEG net (Electrical Geodesics, EGI NA 400 amplifier) against the Cz reference and was sampled at 1000 Hz. Electrooculographic (EOG) signals were measured by 4 facial electrodes located above and on the external side of both eyes. Impedances were kept below 50 kΩ (Ferree et al., 2001). Recorded data were then exported into a custom-made MATLAB format for subsequent preprocessing and analysis. Synchronization with EEG recordings was achieved using Net Station triggers in combination with photodiode markers embedded at predefined visual events, including photographer-name and image onsets. In addition, a square-wave synchronization signal was presented at the onset of each audio-visual movie to verify timing alignment between stimulus presentation and EEG recordings.

### Standardized Measures of Attention

To complement the neural and behavioral measures collected during the experimental sessions, participants completed an additional assessment session focused on standardized measures of attention after their EEG session. During this session, participants completed the Adult Self-Report Scale (ASRS), a standardized self-report questionnaire designed to screen for symptoms associated with ADHD, including difficulties in sustained attention, organization, impulsivity, and distractibility (Kessler et al., 2005). The ASRS consists of 18 items divided into two subscales. Part A includes six items considered most predictive of ADHD symptoms and is commonly used as a primary screening section. Part B includes an additional 12 items assessing broader attentional and behavioral characteristics, such as distractibility, restlessness, and task management. Responses were used to derive Part A scores, Part B scores, and a total attentional symptom score for each participant.

### 2.5 Data Analysis

#### 2.5.1 Behavioral Data Analysis

Behavioral analyses focused on learning outcomes based on the pictures-matrix test and the verbal descriptions provided by the students. For the picture-matrix, accuracy scores were calculated as the percentage of correctly identified photographs for each participant and condition. To assess overall learning across the experiment, pre-test familiarity scores were compared with post-test learning scores using a repeated measures ANOVA (Time X Condition). Since students achieved relatively high scores on the pictures-matrix task, suggesting potential ceiling effects (M = 92.85%, SD = 14.10%), all analyses that examined the connection between neural activity and learning outcomes were conducted with the verbal description scores, which showed greater variability across participants and conditions. For the verbal descriptions part, participants’ responses were scored offline by two double-blinded raters on a 0-3 scale reflecting the level of learning: 0 = no relevant description, 1 = limited or example-specific description, 2 = accurate but concise description of the photographer’s overall style, and 3 = comprehensive and accurate description including key characteristics. To evaluate whether learning modality influenced performance, the mean verbal response scores for each condition were compared using a paired-samples t-test.

##### Attention ratings

Subjective ratings of attention, reported by the students at the end of each block, were analyzed to characterize participants’ engagement during the museum tours. We averaged those scores across conditions and blocks and conducted a paired-samples t-test on these scores to see if there was any difference between conditions.

#### 2.5.2 EEG Data Analysis

##### Preprocessing

EEG data were preprocessed using the FieldTrip toolbox (MATLAB, https://www.fieldtriptoolbox.org, version 20220729). For each participant, the raw EEG data from all trials (both conditions) were appended together for the preprocessing phase. First, all data were band-pass filtered between 0.5 and 40 Hz (4th-order zero-phase Butterworth IIR filter), detrended, and demeaned. Visual inspection was performed to identify and remove gross muscle artifacts, and Independent Component Analysis (ICA) was then used to further remove components associated with horizontal or vertical eye movements and heartbeats. Remaining noisy electrodes, containing extensive high-frequency activity or DC drifts, were removed and interpolated using neighboring electrodes.

##### Attentional modulation of neural speech tracking

As a first step, we examined how active auditory attention toward the teacher modulated the relationship between the teacher’s speech (stimulus; *S*) and the neural activity (response; *R*) relative to the passive hearing condition. To that end, we performed a speech-tracking analysis using TRF, allowing us to compare the coupling between the acoustic features of the speech signal and neural activity across attention conditions. Analysis was conducted using the mTRF MATLAB toolbox (Crosse et al., 2016), and consisted of an encoding and decoding approach.

Each 2.5-minute continuous EEG pre-processed data (corresponding to one block) was segmented into thirteen 10-second trials. The segmented data were then bandpass filtered between 0.5 and 20 Hz using a fourth-order zero-phase Butterworth IIR filter and downsampled to 100 Hz to improve computational efficiency. Then, the neural data were normalized using a z-score transformation, allowing comparability between movies and participants. To extract the acoustic features from the speech signal, a broadband temporal envelope was derived by filtering the audio through a cochlear filterbank, extracting the narrowband envelopes using the Hilbert transform, and averaging across frequency bands. The stimulus signal was then segmented to correspond with the EEG trials. Finally, the resulting envelopes were downsampled to 100 Hz to match the EEG data and normalized using a z-score transformation.

##### Encoding

In the encoding analysis (*forward modeling*), TRFs were estimated separately for each EEG channel using time lags ranging from −150 ms (baseline) to 400 ms relative to the speech stimulus. TRFs were computed using the mTRF toolbox (Crosse et al., 2016), which applies ridge regression (L2 regularization) to improve model generalizability and reduce overfitting. To determine the optimal regularization parameter (λ), we tested a range of λ values between 10⁻² and 10L using a leave-one-out cross-validation procedure. In each iteration, the model was trained on all trials except one and then used to predict the neural response in the held-out trial. For all reported analyses, we selected the λ value that yielded the highest predictive power (Pearson correlation r value) across the largest number of participants. This approach was chosen to maintain comparability of TRF models across participants, rather than optimizing λ separately for each individual participant (see Kaufman & Zion Golumbic, 2023; Levy et al., 2025). To evaluate the goodness of the TRF encoding model with the chosen λ, we assessed whether the predictive power of the model exceeded chance-level performance by generating a null distribution of repeatedly shuffled S and R trials in each condition. This allows us to preserve the overall acoustic features of the signal while disrupting the temporal correspondence between the S and R. The predictive power obtained from the real, non-shuffled data was then compared against this null distribution, showing that for both conditions the TRF performance exceeded chance levels (500 permutations, *p* < .0001). Finally, to identify the significant spatiotemporal differences between attention conditions, statistical cluster analyses (Oostenveld et al., 2011) were conducted using the FieldTrip toolbox. Cluster-based permutation tests were performed using a dependent-samples t-test with Monte Carlo randomization (500 permutations). Spatial clusters were defined based on neighboring electrodes identified using a distance-based approach (alpha < .025), and cluster significance was evaluated using the maximum cluster-level statistic (alpha < .05).

##### Decoding

To complement the encoding analysis, we applied a TRF decoding (*backward modeling*) approach. In contrast to the encoding model, which predicts neural responses from the speech stimulus, the decoding model reconstructs the speech envelope from multichannel neural activity. Reconstruction accuracy was quantified as the predictive power of the model, measured as the Pearson correlation coefficient between the reconstructed speech envelope and the original speech envelope. We used time lags between the S and R ranging from −400 to 0 ms, such that negative lags reflected neural responses following the stimulus. No baseline period was included, as this would reduce decoding performance. Similar to the encoding model, a ridge-regression regularization approach was applied, and reconstruction quality was evaluated using a leave-one-out cross-validation procedure across a range of ridge parameters. The λ value selected for all reported analyses produced the highest predictive power for the largest number of participants. Statistical significance was assessed using a permutation test (100 permutations), in which reconstruction accuracy was compared against a null distribution generated by mismatching speech stimuli and neural responses across trials within each condition. Reconstruction accuracy exceeded chance levels (*p* < .0001). To assess whether reconstruction accuracy differed between attention conditions, statistical comparisons of decoding performance were conducted at the participant level using paired-samples t-tests.

##### Individual differences in attentional modulation and learning

To examine whether individual differences in attentional allocation were related to learning outcomes, we derived an ‘*Attention index*’ for each participant. This index was based on TRF decoding performance and reflected the relative difference in speech reconstruction accuracy between the two task conditions, providing an individualized measure of the participant’s ability to shift attention toward auditory stimuli when needed. To examine whether this ability was specifically related to learning new information from listening, rather than reflecting general comprehension abilities, we conducted a Pearson correlation analysis between the neural ‘*Attention index*’ and students’ verbal response scores across both learning conditions. To avoid including unreliable neural estimates, analyses included only participants who showed above-chance decoding performance relative to a permutation-based null distribution (n = 36).

#### 2.5.3 Teacher-Informed Attention Profiles and Attentional Modulation

A key objective of the current study was to move beyond laboratory-based measures of attention to students’ everyday behavior in the classroom. We therefore examined whether teacher-informed attentional profiles could contextualize students’ neural responses and whether neural measures could, in turn, explain meaningful variation in classroom attention. A separate meeting was conducted between the research team and the teacher after the final experimental session. During this meeting, the teacher was asked to rank all the participating students according to their typical level of attention in the classroom, ranging from relatively high to relatively low. A median split was then conducted to divide all participants into two groups: “High-attention profile” and “Low-attention profile”.

At the time of the session, the teacher had known the 5th-grade students for approximately 5 months, and the 6th-grade students for approximately 15 months; thus, the ranking procedure was conducted separately for 5th- and 6th-grade students to account for differences in teacher familiarity with the classes. To assess the extent to which the teacher-based evaluations aligned with standardized self-reported attentional tendencies, correlation analyses were computed between the teacher’s ratings and students’ scores on the ASRS questionnaire.

No explicit criteria or predefined behavioral features were provided for the ranking process, allowing the teacher to rely on their naturalistic observations and long-term familiarity with the students. Following the ranking procedure, the teacher was asked to verbally describe the main behaviors and characteristics that informed their evaluations. According to the teacher’s descriptions, students classified within the high-attention profile group were characterized by behaviors such as:

● Demonstrating body language consistent with active listening (e.g., looking toward the teacher, reduced side conversations, limited fidgeting).
● Efficiently processing and following instructions without requiring repeated clarification.
● Showing stronger memory and recall for previously discussed information, and independently using organizational tools when needed.
● Displaying higher intrinsic motivation toward learning, including greater care, depth, and engagement in classroom projects and assignments.

##### Group Differences in attentional modulation

To examine if students with different in-class attentional profiles differed in their ability to selectively allocate attention to the teacher’s speech during natural learning, we compared neural speech tracking across task conditions within each group. To allow direct comparison with the analyses conducted in the full cohort, the same TRF encoding and decoding procedures described above were repeated independently within each group. Then, to determine whether these effects were specific to teacher-informed attentional profiles or could instead be explained by objective learning performance, we repeated the same analyses after dividing participants into high- and low-learning groups based on a median split of their verbal learning scores. TRF encoding and decoding analyses were conducted independently within each learning group. Comparing these results with those obtained using teacher-defined attentional profiles allowed us to evaluate whether individual differences in attentional modulation were better explained by objective learning performance or by teachers’ accumulated knowledge of students’ everyday attentional behaviors.

## 3. Results

### 3.1 Behavioral results

To assess learning, we first compared performance on the pre-test and post-test *Pictures-matrix* across both attention conditions combined. A repeated-measures ANOVA (Time × Sensory-modality) revealed a significant main effect of Time, indicating that students demonstrated learning following the learning phase in both conditions [F_(1,38)_ = 278.20, p < .001]. No significant main effect of Condition was observed [F_(1,38_) = 0.31, p = .57], nor was there a significant Time × Condition interaction [F_(1,38)_ = 0.03, p = .86]. Since students achieved relatively high scores on the pictures-matrix task, suggesting potential ceiling effects (M = 92.85%, SD = 14.10%), all further analyses that examined the connection between neural activity and learning outcomes were conducted with the verbal description scores, which showed greater variability across participants and conditions.

Analyses of the *Verbal responses* revealed a condition effect such that participants provided more accurate and detailed descriptions of the photographers’ styles when they listened to their teacher, compared to when they learned by looking [t_(38)_ = 2.76, p < .01]. Interestingly, although students produced higher-quality verbal descriptions in the active listening condition, they rated this condition as less engaging [t_(38)_ = −2.01, p < .001]. A marginally significant effect was also observed for attention ratings, suggesting that students reported lower levels of attention when asked to listen to their teacher, compared to when asked to learn by looking [t_(38)_ = −2.49, p = .052]. A summary of the behavioral results is presented in Figure 3.

**Figure 3:**
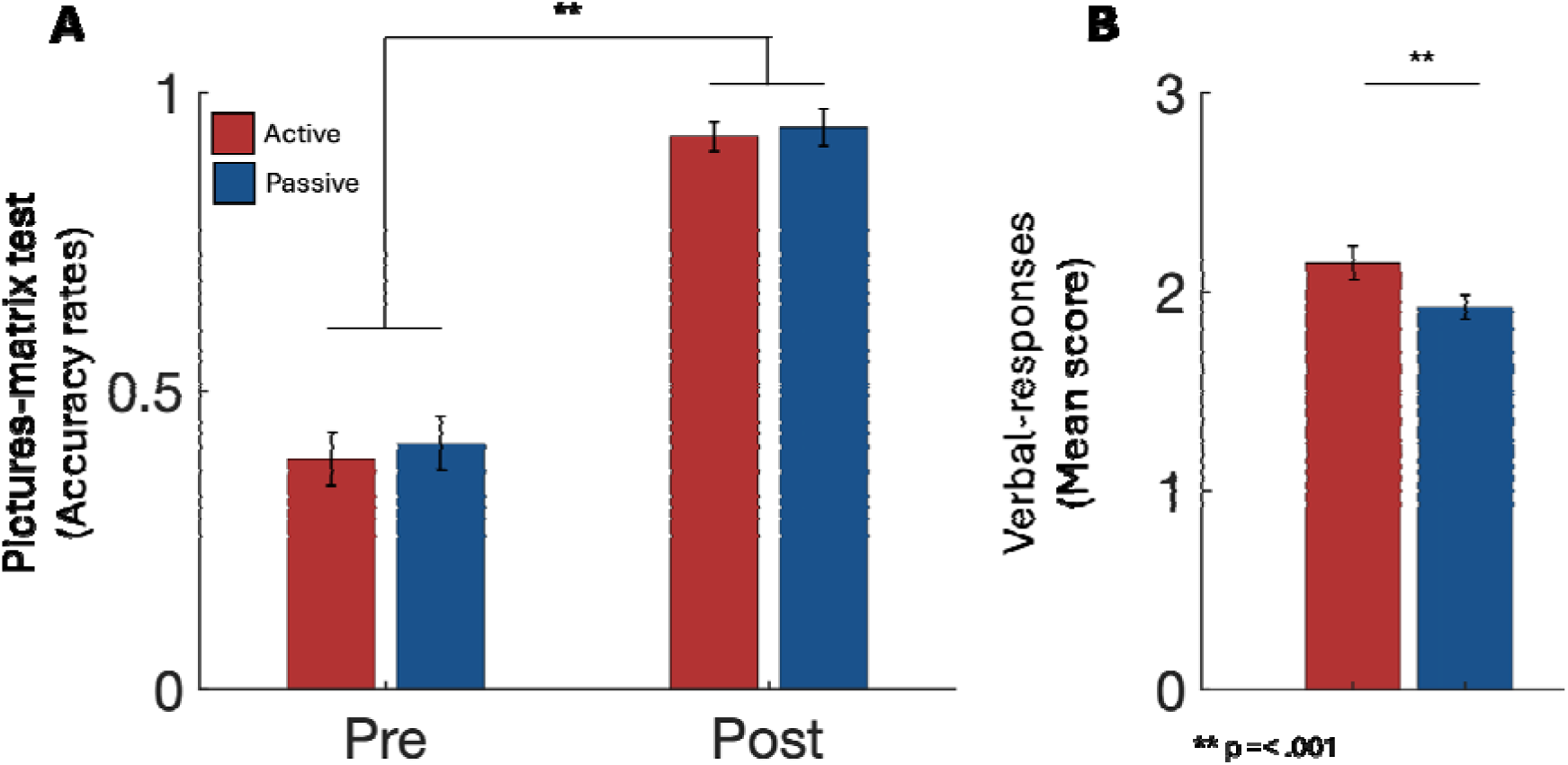
Behavioral learning outcomes following auditory and visual learning. **(A)** Performance on the pictures-matrix test administered before (pre-test) and after (post-test) the learning session. **(B)** Mean verbal-response scores following the learning session. Error bars represent ± SEM.

### 3.2 In the context of education, auditory attention enhances late-stage speech processing

Speech tracking analysis of responses toward the teacher’s speech, in both conditions, yielded robust TRF models (predictive power r = 0.01, p < 0.001 vs. permutations). Visual inspection of the TRFs revealed two positive peaks at ∼ 50 ms and ∼170 ms and one negative peak around 100ms, a canonical three-peak TRF, analogous to the auditory event-related potentials P1-N1–P2 complex (Carta et al., 2023; Lalor et al., 2009).

Figure 4 shows the grand-average encoding TRFs for the two task conditions. Cluster-based permutation statistics corrected for multiple comparisons revealed a significant attentional modulation of the TRF at late latencies, with enhanced neural responses when attention was actively directed toward the teacher’s speech [t_(38)_ = 4.021, p < .05]. The active listening condition showed a significantly larger late positive component between 105–213 ms. Sensor-level topographies further indicated that these differences were primarily driven by activity over frontal electrode sites. No differences between conditions were observed for the earlier positive or negative peaks, suggesting that active listening, compared to passive hearing, primarily modulated late stages of speech processing rather than early sensory encoding. Next, we examined whether whole-channel speech reconstruction captured the attentional modulation observed in the encoding analyses. We compared TRF decoding performance between the active and passive listening conditions using a paired-samples t-test. Results revealed a marginal group-level difference, with higher speech reconstruction accuracy in the active listening condition than in the passive listening condition [t_(38)_ = 1.77, p = .08].

**Figure 4:**
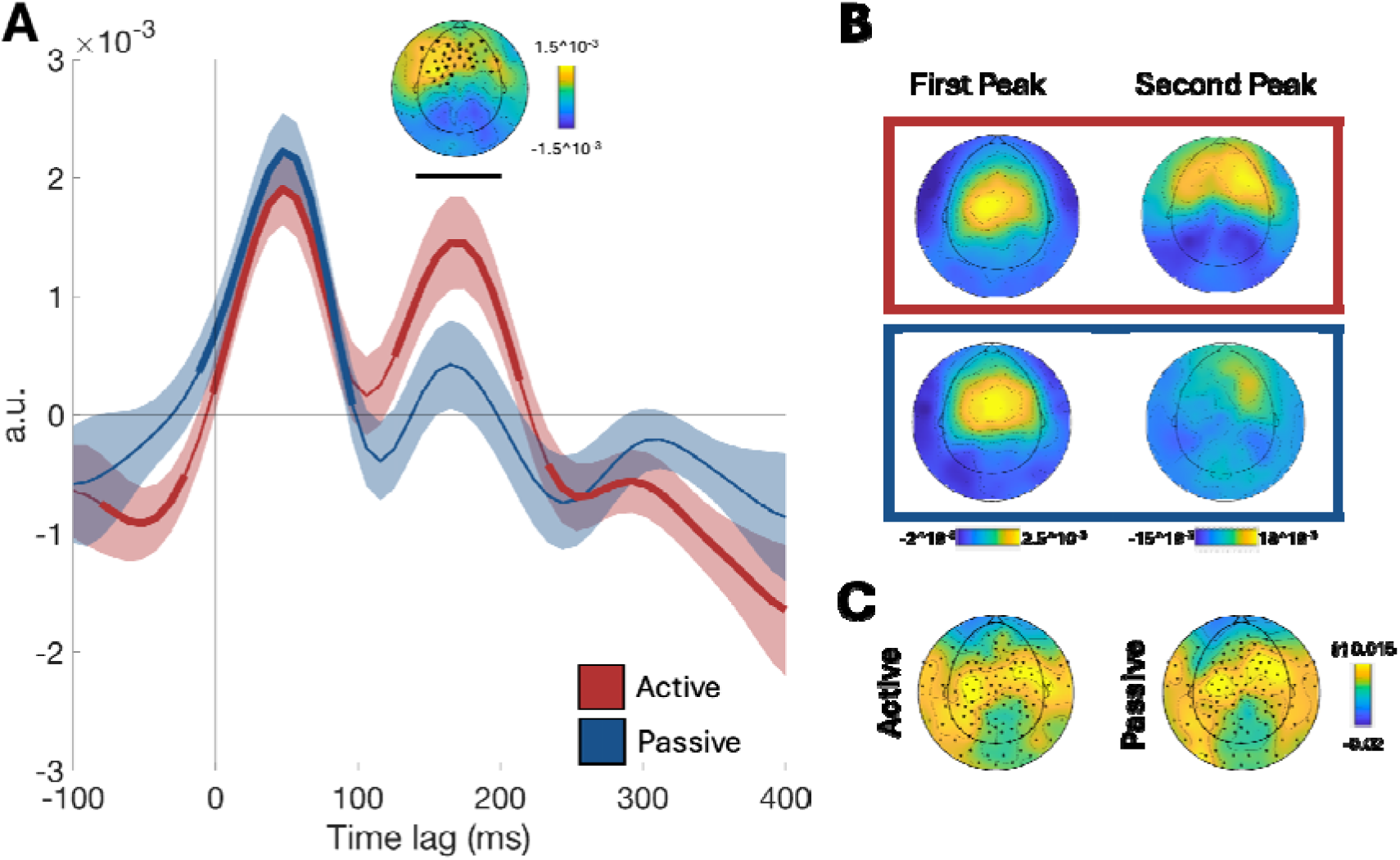
TRF encoding analyses. (A) Grand-average TRFs derived from the encoding model for the active listening (red) and passive hearing (blue) conditions. Shaded areas represent ± SEM across participants. Cluster-based permutation testing revealed a significant difference between conditions at 105–213 ms (black bar), indicating enhanced speech processing when attention was directed toward the teacher’s speech. Topographies represent the spatial distribution of the significant condition effect (Active > Passive Bold segments of the TRF represent temporal clusters where the response significantly differs from zero. Asterisks on the scalp map indicate electrodes contributing to the significant cluster identified by the permutation analysis. (B) Scalp topographies of the encoding model weights at the major TRF peaks for each condition. Topographies are shown for the early positive peak (∼50 ms; left column) and the late positive peak (∼170 ms; right column). The upper row corresponds to the active listening and the lower row to the passive. (C) Topographical distribution of the encoding model predictive power values (Pearson’s r) across scalp electrodes for the active (left) and passive (right) conditions. Asterisks indicate electrodes for which encoding performance exceeded the significance threshold derived from the permutation analysis (95th percentile of the null distribution; r > 0.0024).

### 3.3 Late-Stage Attentional Modulation Selectively Predicts Individual Differences in Learning from Spoken Instruction

To examine whether students’ ability to shift their auditory attention was related to learning outcomes, we conducted a correlation analysis between the ’A*ttention index’* (i.e., the difference in decoding performance between the active and passive conditions) and verbal response scores. Results revealed that participants who exhibited larger increases in decoding performance during active listening produced higher-quality verbal responses when they were asked to learn by listening to their teacher (r = 0.43, p < .001), but not when they learned through visual representations (r = 0.12, p = .47). Meaning that stronger modulation of auditory attention was specifically linked to improved learning from spoken educational information, rather than reflecting a general relationship with overall comprehension or verbal abilities (See Figure 5).

**Figure 5:**
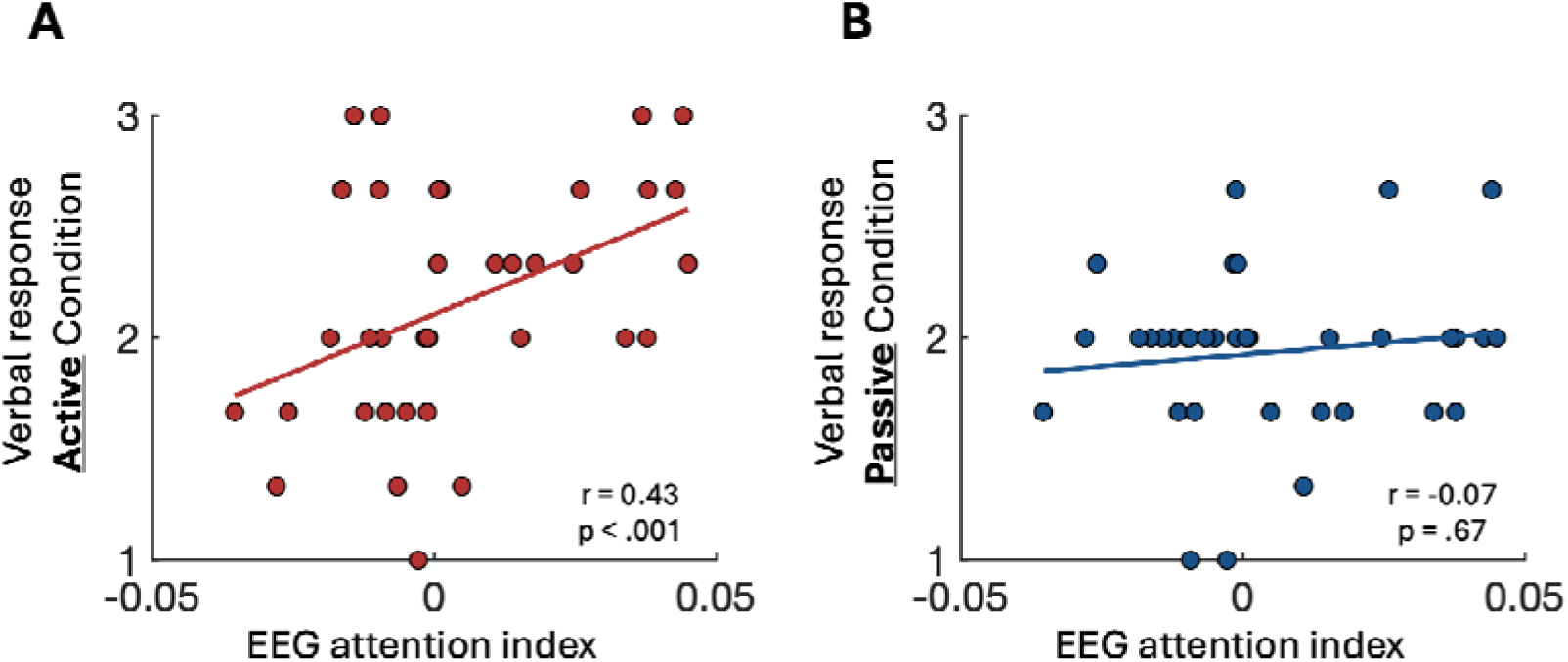
Relationship between attentional modulation of neural speech tracking and learning outcomes. (A) Association between the attention index (defined as the difference in decoding performance between the Active Listening and Passive Hearing conditions) and verbal learning outcomes in the active listening condition. (C) Association between the attention index and verbal learning outcomes in the passive hearing condition.

**Figure 6:**
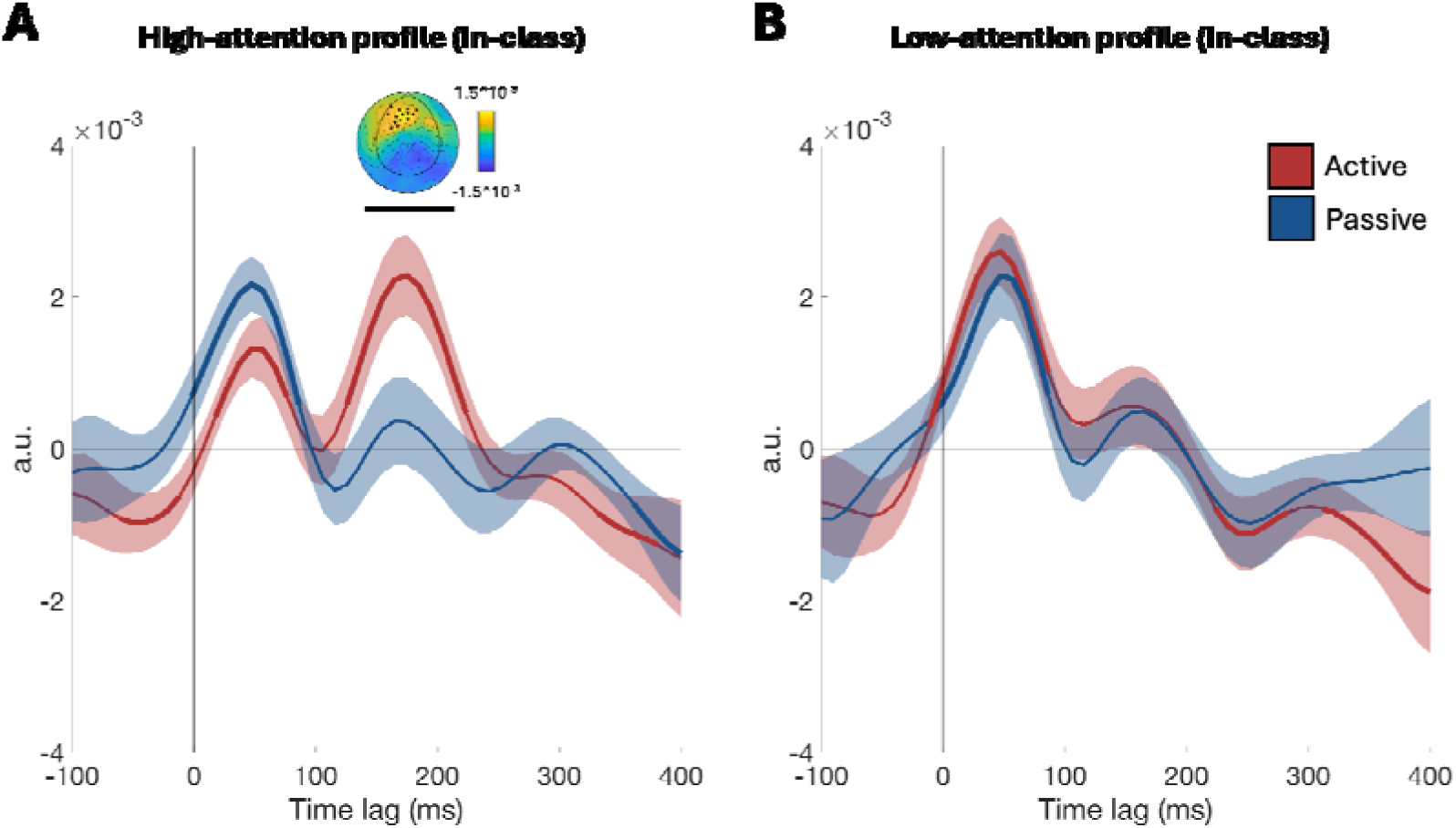
Attention-related modulation of neural speech processing in students with high and low attentional profiles. Grand-averaged TRFs for students with high (left panel) and low (right panel) attentional profiles, comparing the active (red) and passive (blue) conditions. Shaded regions indicate ±1 SEM. The inset topography illustrates the spatial distribution of the attention-related effect. Cluster-based permutation testing revealed a significant difference between conditions for the high-attention profile students only at 136–203 ms (black bar). Topographies represent the spatial distribution of the significant condition effect (Active > Passive). Bold segments represent temporal clusters where the response significantly differs from zero. Asterisks indicate electrodes contributing to the significant cluster identified by the permutation analysis.

### 3.4 Teacher-Informed Classroom Attention Profiles Distinguish Late-Stage Attentional Modulation

To validate the teacher-informed attention profiles, we first examined whether teachers’ ratings were associated with students’ self-reported attentional tendencies measured using the ASRS questionnaire. Results revealed a significant positive correlation between the teacher scores and ASRS Part B scores, indicating that students rated by the teacher as showing greater attentional difficulties also reported higher levels of attentional and behavioral symptoms (r = .43, p = .012). In contrast, the correlation between teacher scores and ASRS Part A, which is usually used for clinical assessments, did not reach significance (r = .28, p = .116).

According to the median teacher-rating score, we divided the group of participants into two groups: a low-attention profile group (n = 18) and a high-attention profile group (n = 21). Figure 5 shows the TRF encoding responses for each group separately. Participants identified as exhibiting a high-attention classroom profile showed significantly greater late-stage (136 and 203 ms) neural speech tracking when attending to the teacher than when attention was directed away from the auditory stream [t_(20)_ = 4.3, p < .05]. In contrast, this pattern was not observed in the low-attention profile group, who showed similar speech-tracking responses across task conditions and less distinct differentiation between active listening and passive hearing (p =.68). For the decoding analysis, a similar pattern emerged, although the effect reached only marginal significance. articipants identified as having a high-attention classroom profile demonstrated a trend toward stronger decoding performance when attention was directed toward the teacher’s speech compared to the visual-attend condition [t_(20)_ = 2.01, p = .058], whereas students in the low-attention profile group showed no significant difference [t_(17)_ = 0.44, p = .66].

Finally, to better understand whether the teachers’ insights contributed information beyond the behavioral learning outcomes alone, we conducted an additional TRF encoding analysis in which participants were divided based solely on their task performance. Unlike the teacher-informed classroom attention profiles, behavioral learning outcomes alone were not sufficient to reliably distinguish neural attentional modulation during speech processing within either of the groups [High performers: t_(18)_ = 3.05, p = .61; Low performers: t_(20)_ = 2.82, p = .60]. No correlation was found between the teacher ranking and the scores of the verbal performance (r = −0.18, p = .29);

## 4. Discussion

In the current study, we sought to capture key features of classroom learning, particularly the need for students to actively regulate their attention to ongoing teacher instruction, and translate these demands into an experimentally tractable paradigm. Our results showed that attention selectively modulated later stages of cortical speech processing, while early sensory encoding remained largely unaffected. Importantly, this neural modulation was meaningfully linked to both learning and classroom behavior: students who showed stronger attentional modulation learned more effectively, and those identified by their teacher as stronger attenders in the classroom exhibited greater neural modulation. Together, these findings reveal a convergence between neural mechanisms of attention, learning outcomes, and teachers’ observations of classroom attention, providing a mechanistic bridge between cognitive neuroscience and real-world learning.

### 4.1 Late-Stage Flexible Resource Allocation Supports Learning from Speech

Understanding how attention supports education represents a critical challenge at the intersection of cognitive neuroscience and real-world learning. Teachers consistently identify students’ ability to engage and regulate their attention as central to successful learning, yet the cognitive and neural mechanisms underlying these everyday manifestations of attention remain poorly understood. Compared with common laboratory tasks involving isolated words or decontextualized sentences, meaningful learning places demands that extend well beyond primary sensory processing. Students must extract meaning, integrate information over time and with prior knowledge, generate and update representations, and encode information in a form that supports subsequent memory and behavior (Mayer, 2014, 2024; Vygotsky, 2011). These operations place substantial demands on cognitive resources even when the incoming information itself is clear and readily perceptible. The present study was designed to capture these demands: participants listened to natural, continuous instruction delivered by their own teacher, engaged with materials aligned with their classroom curriculum, and simultaneously managed competing, learning-relevant visual information.

This distinction between the perceptual demands of hearing speech and the cognitive demands of learning from speech is particularly important for interpreting our neural findings. Speech is a highly salient and behaviorally relevant auditory signal that is readily tracked even when attentional resources are directed elsewhere (Liberman & Whalen, 2000; Zoefel & Kösem, 2024). When speech is clear and intelligible, cortical speech tracking can persist while attention is engaged elsewhere (Kong et al., 2014; Lee et al., 2016; Vanthornhout et al., 2019; Wild et al., 2012). By contrast, when listening becomes perceptually demanding because of competing speakers, background noise, or acoustic degradation, attentional modulation becomes more pronounced (Brodbeck et al., 2020; Crosse et al., 2016; Davis & Johnsrude, 2003; Di Liberto et al., 2015; Ding & Simon, 2012; Fiedler et al., 2019; Puvvada & Simon, 2017).

Here, however, late-stage attentional modulation emerged despite favorable listening conditions. Participants listened to their own teacher; speech was clear, age-appropriate, and adapted to their learning level. This pattern suggests that the attentional demands observed here arose not from the perceptual challenge of encoding speech, as reflected by preserved early temporal encoding across attentional states, but from the cognitive demands of learning from what was heard. In other words, even when speech is readily encoded, transforming that signal into information that can be understood, integrated, and learned may require additional attentional resources.

This interpretation is further supported by evidence that speech processing is fundamentally shaped by the broader context in which communication and learning occur. Speaker familiarity, prior knowledge, and the pedagogical structure of learning materials can all shape the neural processing of continuous speech by modulating listening effort, with particularly pronounced effects at later stages of processing (Baxter et al., 2025; Broderick et al., 2019; Har-shai Yahav et al., 2024; Johnsrude et al., 2013; Piazza et al., 2025). Altogether, the predominance of late-stage attentional modulation we observed may not reflect the need to resolve perceptual competition, but rather the cognitive demands of navigating a rich and dynamically unfolding learning context. Considering the temporal and spatial characteristics of the effect observed here, we suggest that meeting these demands relies on the recruitment of higher-order, top-down processes that support the selection, integration, and maintenance of learning-relevant information (Cavanagh & Frank, 2014).

The importance of efficiently allocating limited cognitive resources becomes particularly salient when considering the developmental stage of the learners themselves. Although adolescents generally demonstrate adult-like selective attention under many laboratory conditions, the perceptual and cognitive capacity that supports these mechanisms continues to mature into late adolescence (Hobbiss & Lavie, 2024; Huang-Pollock et al., 2002). Hobbiss et al. (2024) established that selective attention per se, the foundational ability to completely suppress irrelevant sensory distractors, is fully mature by early adolescence. However, they found a distinct developmental trajectory regarding how perceptual load modulates this selection: Younger adolescents had a reduced overall perceptual and cognitive capacity, causing attentional resources to reach capacity under lower task demands than in older adolescents or adults.

The present findings identify late-stage attentional modulation as a neural mechanism that may be particularly important for supporting learning during the school-age years, when cognitive and attentional systems are still developing. Children’s more limited processing capacity may leave fewer resources available to accommodate the additional demands imposed by complex learning environments. Consistent with this account, speech-in-noise studies show that school-age children require higher signal-to-noise ratios than adults to achieve comparable speech recognition and comprehension (Bonino et al., 2021; Leibold, 2017). By demonstrating flexible attentional modulation of speech processing during learning, our findings extend the developmental literature on neural speech processing beyond its predominant focus on infancy and early childhood (Araújo et al., 2024; Attaheri et al., 2022; Di Liberto et al., 2023; Kalashnikova et al., 2018; Kojima et al., 2026; Pérez-Navarro et al., 2024; Van Hirtum et al., 2023), highlighting the continued role of attentional mechanisms as children navigate the increasingly complex cognitive demands of classroom learning.

### 4.2 Toward an Ecological Understanding of Attention

The field of educational neuroscience is reaching a critical consensus: understanding cognitive functions requires naturalistic paradigms that capture real-life learning without sacrificing empirical rigor. The framework presented here underscores that authentic research-practice partnerships between neuroscientists and educators do more than just improve the ecological validity of experimental tasks - they demonstrate how educators’ expertise can be integrated into experimental research, even within a controlled laboratory setting, to provide novel insights that help bridge theoretical constructs and the realities of classroom learning.

Over the past decade, attention research has increasingly moved toward more naturalistic experimental contexts, adopting paradigms that incorporate natural speech and virtual environments (Levy, Korisky, et al., 2025; Levy, Libman Hackmon, et al., 2025; Mangalmurti et al., 2020; Parsons et al., 2017; Rizzo et al., 2000; Seesjärvi et al., 2022). Yet, increasing the ecological validity of a paradigm does not necessarily mean that it captures the specific cognitive and attentional demands of learning. Drawing on the broader RPP literature, which emphasizes the value of integrating partners with firsthand, system-level expertise into the research process, we argue that involving educators in paradigm development can inform researchers which features of real-world learning are most meaningful to preserve experimentally and thus shape research questions, experimental paradigms, and the interpretation of finding (Coburn & Stein, 2010; Farrell et al., 2022; Gosavi et al., 2026; McGeown & Sjölund, 2026; Toomarian et al., 2024; van Atteveldt et al., 2020; Wilcox et al., 2021). Moving toward a collaborative model also enables the integration of complementary sources of evidence across timescales and contexts. While neuroscientific protocols typically capture only a brief snapshot of a student as an experimental participant, teachers possess a longitudinal perspective developed over an entire school year, allowing neural measures to be interpreted within the broader context of students’ everyday learning and behavior. This deep familiarity with classroom dynamics and individual learning trajectories offers a vital interpretive lens for neuroscientists, allowing us to identify meaningful sources of cognitive variability that would otherwise be dismissed as statistical noise.

The value of this collaborative approach also allowed us to test whether neural mechanisms identified under controlled experimental conditions mapped onto students’ everyday attentional behavior. Working with an experienced classroom teacher, we systematically co-designed an attention paradigm that combined established laboratory methods with a learning context reflecting meaningful features of classroom instruction. Importantly, the teacher’s role extended beyond paradigm development. Teacher observations provided information beyond task performance alone, offering an independent, ecologically grounded measure against which individual differences in neural attention could be evaluated. The convergence between teacher-informed attention profiles and experimentally identified late-stage attentional modulation suggests that this neural mechanism captures meaningful variation in attention that extends beyond the laboratory context. More broadly, this approach illustrates one way to bridge experimental cognitive neuroscience and educational practice. Genuine co-creation remains relatively rare, in part because researchers and educational partners often operate with different professional languages, timelines, priorities, and definitions of success (Ansari & Coch, 2006; Brookman-Byrne & Commissar, 2019; van Atteveldt et al., 2018, 2020). Meaningful co-design also requires sustained collaboration, mutual engagement, and substantial investment from both researchers and educational partners k(Kelly, 2004). Although such an approach may not be feasible or necessary for every research question, when these conditions can be established, collaborative design can support experimental paradigms that are both theoretically precise and grounded in educational practice. Ultimately, such partnerships may help move the field beyond making laboratory paradigms more naturalistic toward identifying which neural mechanisms of attention are genuinely relevant to learning in the classroom.

### 4.3 Limitations

Several limitations should be considered when interpreting the present findings. First, although the paradigm was designed to capture key features of authentic classroom learning, neural activity was still recorded under controlled laboratory conditions. An important next step will therefore be to examine whether the late-stage attentional mechanisms identified here similarly emerge and fluctuate during ongoing instruction in the classroom. Second, the present study focused on a single learning context and set of instructional materials. Including control conditions that systematically vary learning content, task demands, or instructional context would help determine whether the observed late-stage modulation reflects a general mechanism supporting learning or is specific to the demands of the current paradigm. Finally, the findings should be replicated in larger and more diverse student populations, across different ages, educational settings, and learning profiles, to establish the robustness and generalizability of the proposed mechanism.

### 4.4 Conclusion

In the current study, we show that learning from speech depends not simply on whether information reaches the brain, but on how flexibly neural processing is adjusted to meet the goals of the learner. Even when a teacher’s speech was clear and robustly encoded, attention selectively modulated later stages of cortical processing, and students who more effectively adjusted this processing across learning demands were also those who learned more successfully and were recognized by their teachers as stronger attenders in the classroom. These findings position late-stage attentional modulation as a candidate mechanism through which limited cognitive resources are flexibly allocated during learning. More broadly, they demonstrate what becomes possible when cognitive neuroscience moves beyond asking whether laboratory-defined mechanisms generalize to education and instead allows educational contexts to inform which mechanisms we seek to understand. By bringing experimental precision into dialogue with the longitudinal expertise of educators, we can begin to build a science of attention that captures not only how the brain selects information, but how learners use their limited cognitive resources to transform that information into knowledge. Such an approach offers a path toward an educational neuroscience in which classrooms are not merely settings in which theories are tested, but partners in shaping the theories themselves.

