## Supplemental Table 1 for "Bringing Attention to Education: Revisiting the Neural Mechanisms of Selective Attention in Naturalistic Learning"

**Table S1: Demographic and Clinical Characteristics of the Sample**

A total of 39 participants were included in the final analysis of the first (age range: 10.2–12.3 years, M = 11.31, SD 0.60. The sample comprised 18 males (46.1%) and 21 females (53.8%). Most participants were right-handed (94.8%), and had English as their primary language (79.5%), while 8 participants (20.5%) reported being bilingual (English and one other language). In terms of ethnicity, 16 (41%) identified as White, 12 (30.7%) as Asian, and 1 (2.5%) as American Indian. A prior diagnosis of attention-deficit/hyperactivity disorder (ADHD) was reported by 5 (12.8%) participants, while 4 (10.2%) indicated a specific learning difficulty, such as dyslexia. Regarding medication, 6 (13.9%) participants were currently taking prescribed medication as detailed in the table.

| **Variable** | **Category** | **Experiment 1 (n=39)** | |
| --- | --- | --- | --- |
|  |  | **n** | **%** |
| Gender | Male | 18 | 46.1 |
|  | Female | 21 | 53.8 |
| Ethnicity | White | 16 | 41 |
|  | Asian | 12 | 30 |
|  | American Indian | 1 | 2.5 |
|  | Multicultural | 10 | 25 |
| Handedness | Right | 37 | 94.8 |
|  | Left | 2 | 5.2 |
| Primary Language | English | 31 | 79.5 |
|  | Bilingual | 8 | 20.5 |
| Former Diagnosis | ADHD | 5 | 12.8 |
|  | Other learning difficulties | 4 | 10.2 |
| Medication Type | Stimulant | 2 | 5.1 |
|  | Antihistamine | 2 | 5.1 |
|  | Anti-inflammatory | 1 | 2.5 |
|  | Mood stabilizers | 0 | 0 |
|  | Other | 1 | 2.5 |
|  | No medication use | 33 | 84.6 |

*Values represent counts (n) and percentages (%). ADHD = attention-deficit/hyperactivity disorder.*
